# Molecular Clock and Phylogeny of *Anopheles* species of the subgenus *Nyssorhynchus* (Diptera: Culicidae)

**DOI:** 10.1101/262147

**Authors:** Richard Hoyos-López

## Abstract

The main Phylogenetic hypothesis supporting the Myzorhynchella section as a natural group, but the sections Albimanus and Argyritarsis, do not present clearly resolved relationships, nor is it possible to recover the monophyly of both sections, even within these sections of *Nyssorhynchus*; it has not been possible identify the relationships between the species that make up these taxonomic subdivisions (Sallum, 2000, Sallum, 2002, Bourke, 2011, Foster, 2013). This lack of resolution has been attributed to the effect of few species for phylogenetic studies, making difficult the determination of monophyly of many groups, subgroups and complexes within sections Albimanus and Argyritarsis

We infer the phylogeny of the subgenus *Nyssorhynchus* through the sequences characterized for the molecular markers ND6, COI-Barcode, White and CAD, in addition we calculate the times of divergence for the main lineages corresponding to the sections Albimanus, Argyritarsis and Myzorhynchella using Bayesian approaches.

## 1. Introduction

Malaria is among the most important diseases worldwide with approximately 216 million cases registered for 2011 and an estimated 655,000 deaths in 2010 (Sachs, 2002, WHO, 2011). This disease constitutes an important obstacle for the economic and social development of the affected countries, despite the control programs that have helped to reduce its incidence since 2000, it has been observed that the maintenance of effectiveness in the control programs has decayed and consequently allowed historical re-emergence in endemic regions and rural areas of South-America (Cohen, 2012).

The identification of species is essential for vector incrimination, the measurement of epidemiological risk and the development of management programs for malaria (Sinka, 2010); the focus of these works are guided to mosquitoes belonging to the genus *Anopheles*, which contains all known malaria vectors, with a significant number of representatives considered complex species (Foster, 2013), these are considered morphologically indistinguishable and in other cases phenotypic plasticity makes identification difficult, due to similarity with closely related species (Sallum, 2000). In these cases of taxonomic complexity, only approaches with markers and molecular tools allow to resolve the identity, relationships between species and vector incrimination (Porter & Collins, 1991, Paskewitz, 1993, Cornel, 1996, Cywinska, 2006, Marelli, 2006, Zapata, 2007, Hoyos-López et al. 2012a, Hoyos-López et al. 2012b, Hoyos et al. 2015a, Hoyos et al. 2015b, Hoyos et al. 2015c, Hoyos et al. 2016, Hoyos et al. 2017).

Mosquitoes of the subfamily Anophelinae include 465 formally recognized and 50 undescribed species belonging to species complexes (Harbach, 2007, Foster, 2013), all of them divided into three genera: *Anopheles, Bironella* and *Chagasia*.

*Anopheles* is subdivided into seven subgenres: *Anopheles, Baimaia, Cellia, Kerteszia, Lophopodomyia, Nyssorhynchus* and *Stethomyia*. Nine species of the genus *Anopheles* are considered primary vectors in America, three species belong to the subgenus *Anopheles* and six to *Nyssorhynchus* (Sinka, 2010), however, the latter encompasses at least twelve species considered secondary vectors because of their local and / or regional importance (Bourke, 2011). The identification and separation of anophelines has involved research in phylogenetics and phylogeography, mainly in species of recognized epidemiological importance (Porter & Collins, 1991, Paskewitz, 1993, Cornel, 1996, Sallum, 2000, Cywinska, 2006, Marelli, 2006, Zapata, 2007; Bourke, 2011; Foster, 2013).

The subgenus *Nyssorhynchus* presents a neotropical distribution, including 39 formally recognized species and around six not described but differentiable by molecular markers, subdivided into three sections: Albimanus, Argyritarsis and Myzorhynchella. Sallum et al. (2000) using morphological characters and a wide range of species, is able to determine the monophyly of the subfamily Anophelinae and the subgenus *Cellia, Kerteszia* and *Nyssorhynchus*, which was subsequently corroborated with evidence from nuclear and mitochondrial gene sequences (Krzywinsky 2011; Sallum 2002; Harbach 2007), however, the phylogenetic relationships within the subgenus *Nyssorhynchus* were not clarified. Bourke et al. (2010) and Foster et al. (2013), using DNA sequences from the ND6, White, COI and CAD markers, respectively, provides evidence of statistical support for some lineages, but clear genealogical discrepancies of inferred Bayesian trees are observed for each marker.

The concatenated Bayesian topologies allow to support the Myzorhynchella section as a natural group, but the sections Albimanus and Argyritarsis, do not present clearly resolved relationships, nor is it possible to recover the monophyly of both sections, even within these sections of *Nyssorhynchus*; it has not been possible identify the relationships between the species that make up these taxonomic subdivisions (Sallum, 2000, Sallum, 2002, Bourke, 2011, Foster, 2013). This lack of resolution has been attributed to the effect of few species for phylogenetic studies, making difficult the determination of monophyly of many groups, subgroups and complexes within sections Albimanus and Argyritarsis, also to the effect of low phylogenetic signal for ND6 genes + White used by Bourke et al. (2010) and CAD + COI-Barcode by Foster et al. (2013). Reidenbach et al. (2009), estimates the first approach to the time of divergence between two species of the genus Anopheles, belonging to the subgenera *Stethomyia* (*Anopheles gambiae*) and *Cellia* (*Anopheles atroparvus*), inferring a separation of these taxa about 20–24 million years ago.

Currently, it is considered that a large number of sequences from nuclear and mitochondrial genes can solve phylogenetic problems and infer divergence times like the study by Regier & Zwick (2008), who used sequences of 62 nuclear genes coding for proteins and were able to infer the phylogeny of the phylum Arthropoda and the work of Reidenbach (2009), characterizing seven molecular markers to infer the phylogeny of the subfamilies, tribes and genera representative of the Culicidae family.

In this research, we infer the phylogeny of the subgenus *Nyssorhynchus* through the sequences characterized for the molecular markers ND6, COI-Barcode, White and CAD, in addition we calculate the times of divergence for the main lineages corresponding to the sections Albimanus, Argyritarsis and Myzorhynchella using Bayesian approaches.

## Material and Methods

### Preparation of sequences and alignments

Through Genbank it was possible to obtain the sequences belonging to the ND6 and White markers described by Bourke (2011), and those used by Foster (2013): CAD and COI-Barcode. Manually aligning sets of 40 sequences for each marker in 22 species of the subgenus Nyssorhynchus and two external groups (Anopheles, Stethomyia) (Table 1). Species with little representation were eliminated and emphasis was placed on those that presented information for all the genes used. In this way, individual and one concatenated alignments were constructed (Table 1). For the White and CAD nuclear genes, only blocks with clear positional homology were used, discarding gaps and blocks with multiple ambiguous sites. In COI-Barcode and ND6, amino acid translation was performed to avoid nuclear sequences of mitochondrial origin (NUMTs), pseudogenes and unedited sequences of Genbank. The sequences for each marker were manually evaluated and aligned with Clustalw in Bioedit v7.0.2 (Hall, 2007) and MUSCLE in Mega v5.0 (Tamura, 2011).

**Table 1.**
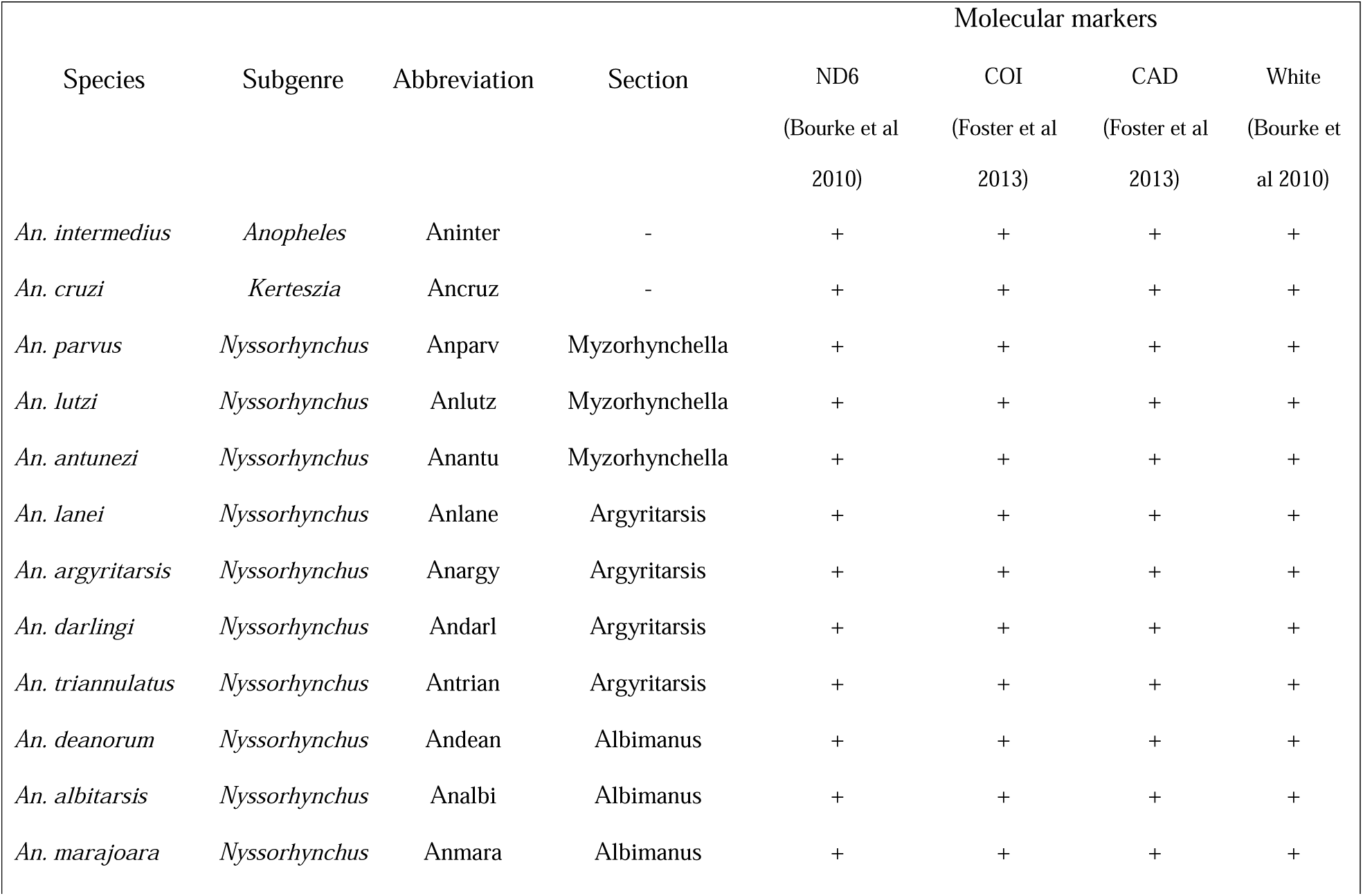
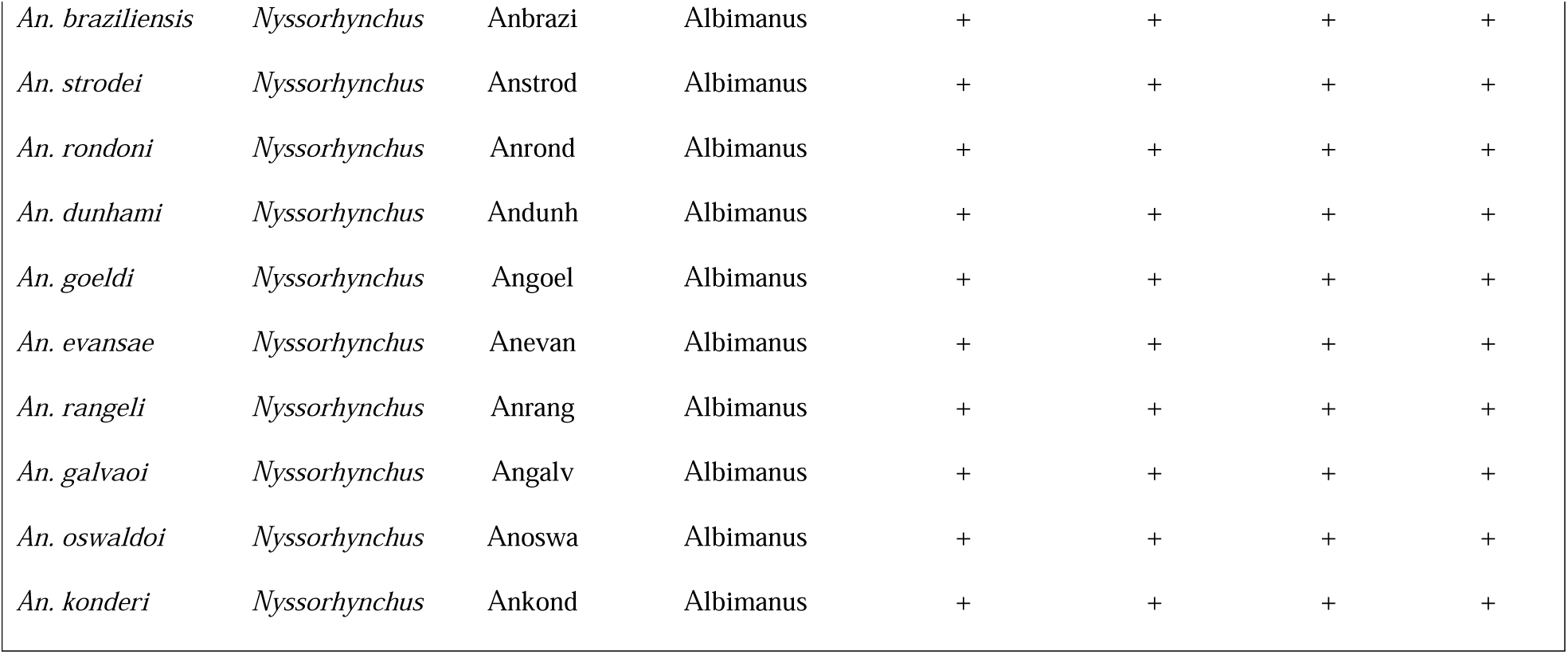
List of species, abbreviations and markers used for phylogenetic inference.

### Estimation of Evolutionary Model

The fasta files for each molecular marker were used to evaluate evolutionary models under the Akaike information criterion (AIC) and likelihood evaluation in the software jModeltestv2.0 (Posada, 2012), starting with an inferred tree in phyml and 24 models for the evaluation (likelihood scores).

### Phylogenetic analysis

The inference by Bayesian methods was applied to sequences of ND6, COI-Barcode, CAD, White and concatenated data using partitioning schemes for previously determined codon positions and evolutionary models. All the analyzes were executed in MrBayesv3.2 (Ronquist, 2012) and each analysis consisted of two simultaneous chains, which were repeated to confirm the convergence of the posterior probability distribution for the parameters estimated by TRACER (Rambaut, 2007).

For the individual gene matrices, each run was 5 million generations and the first 2.5 million were discarded as “burn-in”. The concatenated matrix was analyzed in 20 million generations and the first 10 million generations were discarded. The consensus trees were visualized in TREEVIEW (Page, 1996) and Adobe Illustrator.

### Estimation of divergence times

The inference of separation times for the species of the subgenus *Nyssorhynchus* of *Anopheles* was calculated using a relaxed Bayesian molecular clock (uncorrelated & log-normal) and a “Yule-Calibrated” speciation process (Hoyos et al, 2015a; Hoyos et al. 2015b). Topology of the tree, minimum and maximum intervals of the age of the nodes, and age restrictions by secondary and fossil data were used to define the priors. Reidenbach et al. (2009) determines the age of the lineage of the genus Anopheles in 43.11 million years (age range of nodes: 27.20 – 63.95 million years) by estimating the molecular clock with six nuclear genes. The subgenus Nyssorhynchus was restricted in the minimum age of the node using fossil information (Anopheles dominicanus - 25 million years) (Zavortink, 2000). The XML file for analysis was modeled in BEAUtiv2.0, and the estimate was executed in BEASTv1.8 (Drummond, 2007) and a Markov-Monte Carlo chain of 10 million generations. For the construction of the consensus, 50% of the samples were discarded and with a posterior probability of a minimum clade of 0.5 in the tool in TreeAnnotator v1.8. The consensus tree was visualized in FigTree v1.4.

## Results

The alignments for each marker presented different lengths: CAD (634 nt), White (460 nt), COI-Barcode (606 nt), ND6 (525 nt) and the concatenated matrix presented a length of 2225 nt. For individual markers and the concatenated matrix, the evolutionary model obtained under the Akaike criterion in jMODELTESTv2.0 was GTR + G + I.

The Bayesian trees inferred for the genes ND6 (Figure 1) and White (Figure 4) showed low capacity to resolve the relationships between groups of species and sections within Nyssorhynchus, as was possible to observe with the COI-Barcode markers (Figure 2) and CAD (Figure 3), however, the presence of polytomies and relationships with low inter-species support was observed. The Bayesian tree obtained from the concatenated matrix (Figure 5), allowed to clearly infer the evolutionary relationships between groups and sections within the subgenus, with the exception of *An. arvus* that is located in an ancestral position and external to *Nyssorhynchus*; most later clade probabilities for inferred topology support relationships with high values (0.9–1.0). The species that make up the Argyritarsis section do not constitute a monophyletic evolutionary group and branch lengths with high divergences are observed, and even *An. darling*i is located as an ancestral group and related to *An. parvus*.

**Fig. 1.**
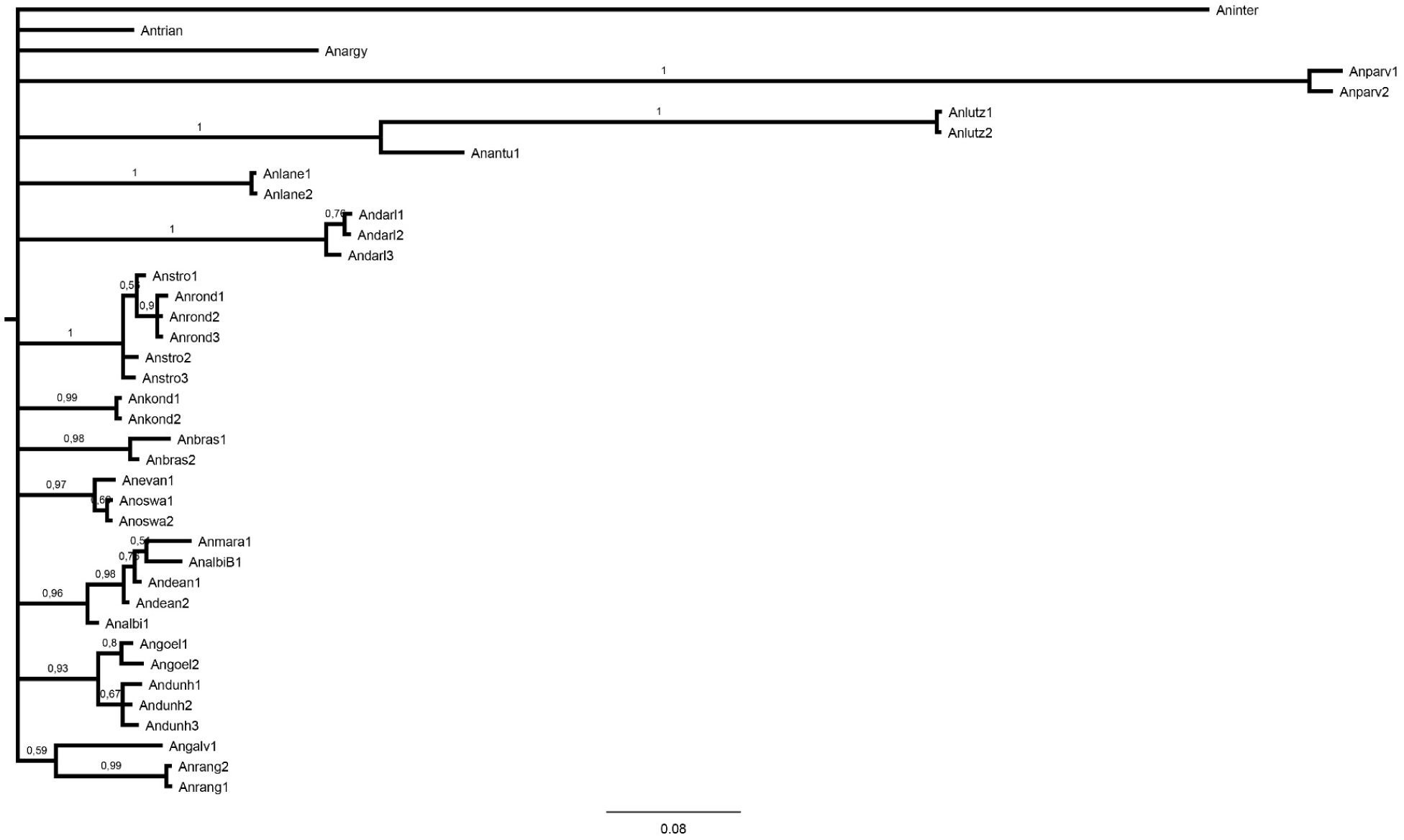
Bayesian consensus tree obtained by 40 sequences of ND6 and 20 species of the subgenus *Nyssorhynchus*. The numbers on the nodes represent the posterior probabilities of clade and the names of the terminals are abbreviated (Table 1).

**Fig. 2.**
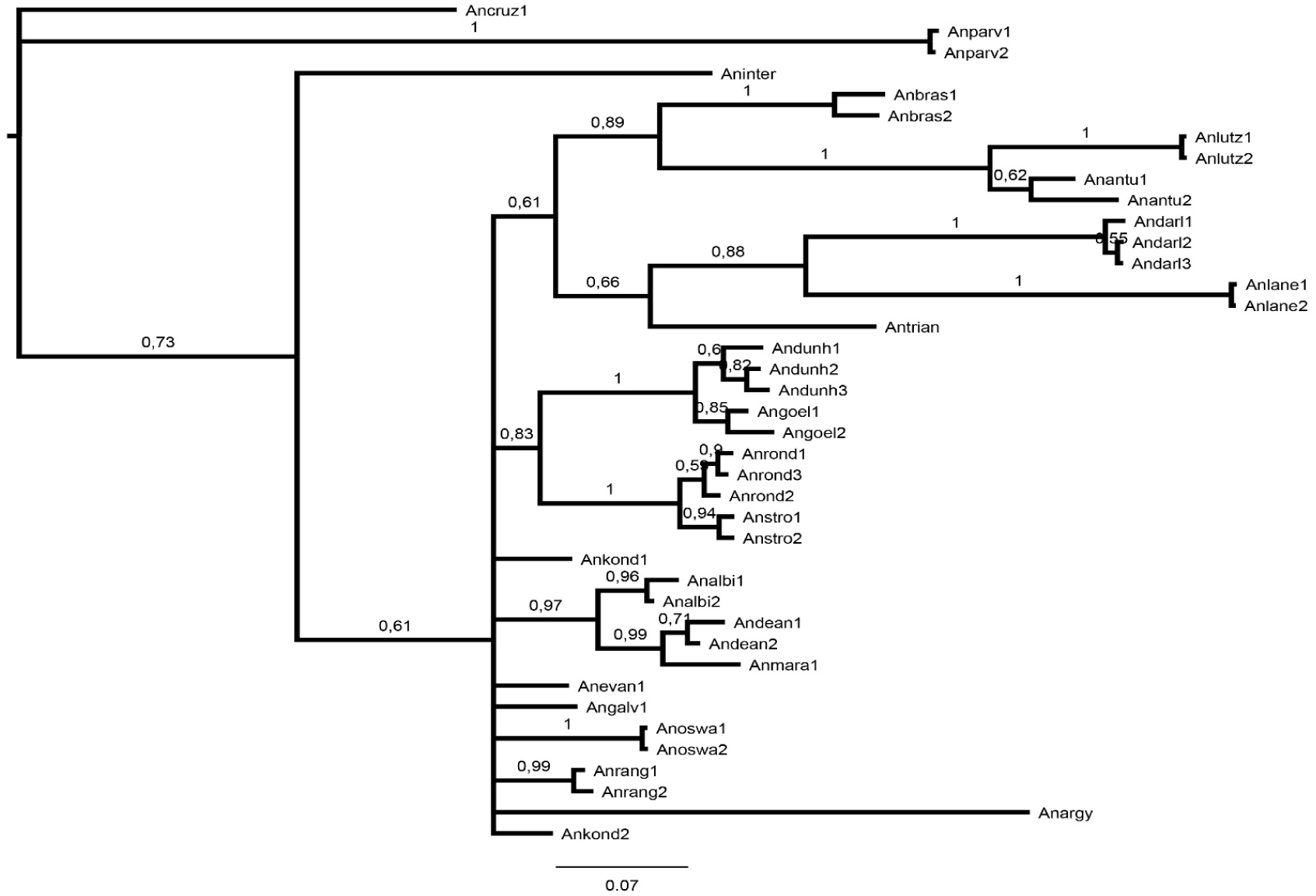
Bayesian consensus tree obtained through 40 COI sequences and 20 species of the subgenus *Nyssorhynchus*. The numbers on the nodes represent the posterior probabilities of clade and the names of the terminals are abbreviated (Table 1).

**Fig. 3.**
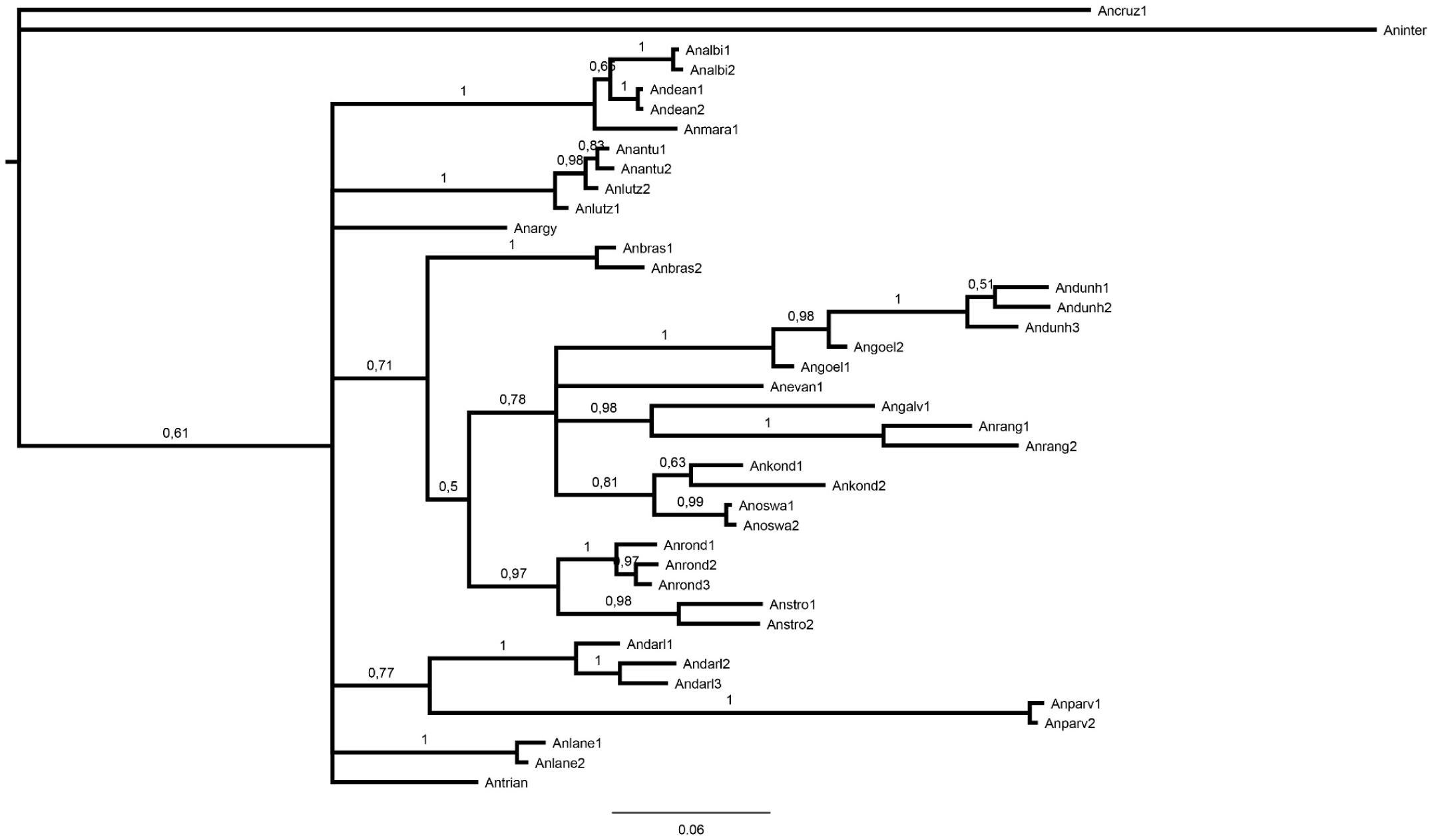
Bayesian consensus tree obtained by 40 CAD sequences and 20 species of the subgenus *Nyssorhynchus*. The numbers on the nodes represent the posterior probabilities of clade and the names of the terminals are abbreviated (Table 1).

**Fig. 4.**
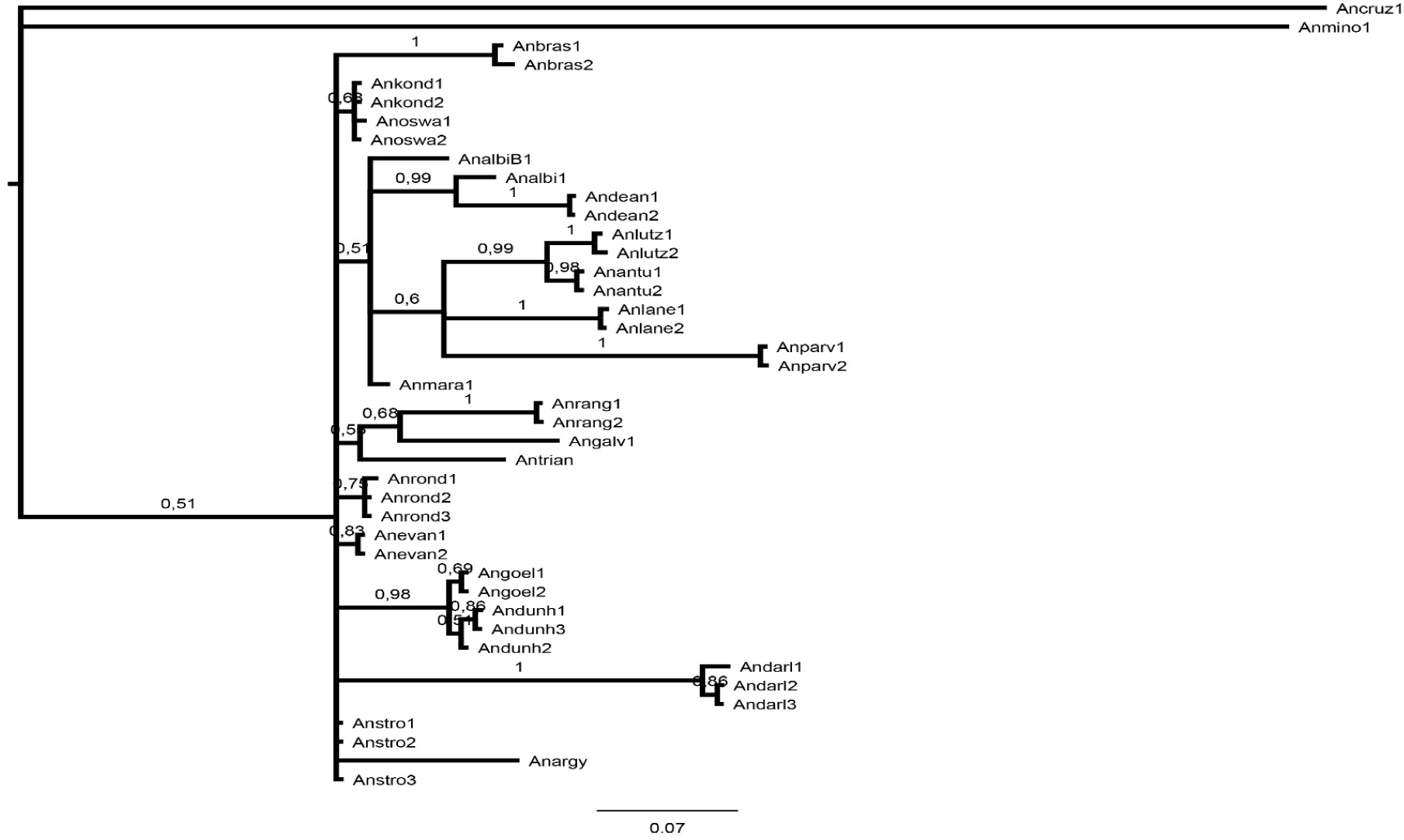
Bayesian consensus tree obtained by 40 White sequences and 20 species of the subgenus *Nyssorhynchus*. The numbers on the nodes represent the posterior probabilities of clade and the names of the terminals are abbreviated (Table 1).

**Fig. 5.**
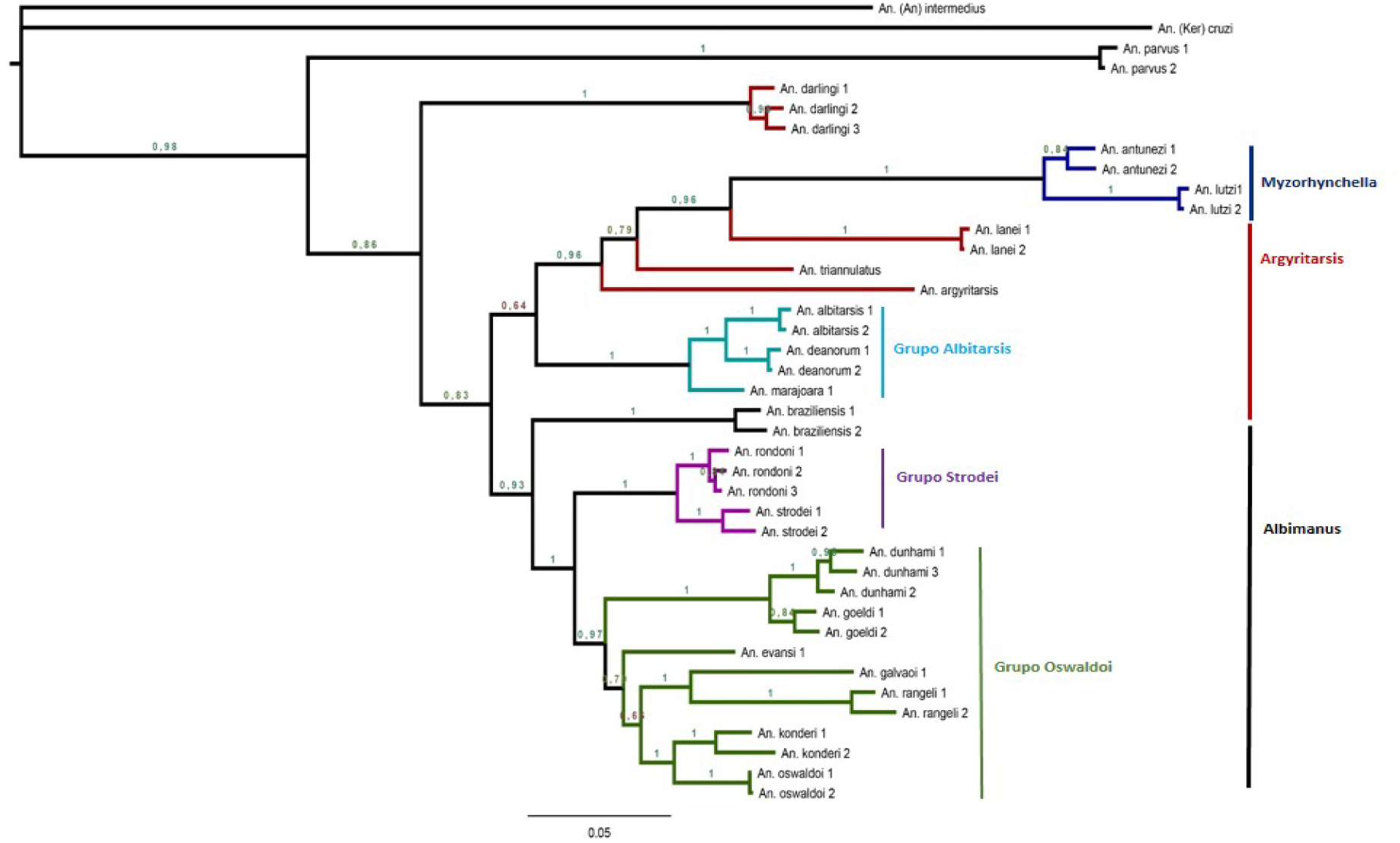
Bayesian consensus tree obtained through the concatenation of 160 sequences of ND6, COI, CAD and White, for 20 species of the subgenus *Nyssorhynchus*. The numbers on the nodes represent the posterior probabilities of clade and the bar on the lower part (0.05), indicates the number of expected substitutions in the estimated branch lengths.

**Fig. 6.**
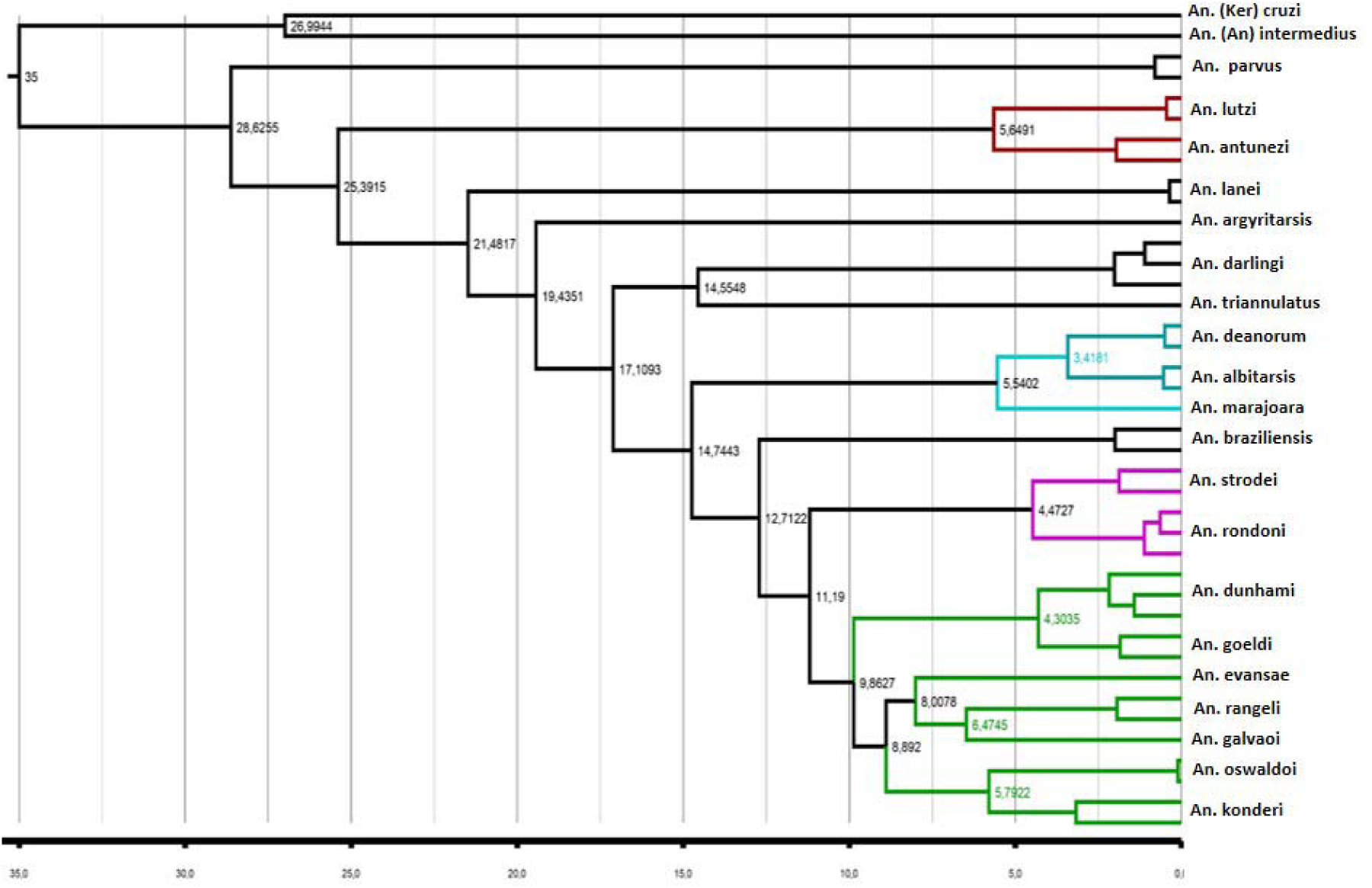
Estimates of divergence obtained by Bayesian inference (Uncorrelated molecular clock, Calibrated Yule) and the concatenation of 160 sequences of ND6, COI, CAD and White, for 20 species of the subgenus *Nyssorhynchus*. The numbers on the nodes represent the estimated ages for the phylogenetically reconstructed lineages.

The Myzorhynchella section conformed in this analysis by *An. antunezi* and *An. lutzii*, presents evident relations with *An. lanei, An. triannulatus* and *An. argyritarsis*, all considered within the Argyritarsis section. The affinity of the Albitarsis group is also observed, to the clade formed by Myzorhynchela + Argyritarsis.

The Albimanus section has closely related Strodei and Oswaldoi groups, with *An. braziliensis* as the species most external to the clade formed by both groups.

The estimation of divergence times shows the formation of the main lineages of *Nyssorhynchus* in the range of 21.48–14.74 million years corresponding to the Miocene (23.03–7.24 ma) and the species located in the groups Albitarsis, Strodei and Oswaldoi seem to diverge in the Pliocene (5.3–3.6 ma) -Pleistocene (2.5 – 0.16 ma).

## Discusión

The estimation of evolutionary relationships from individual markers has proved to be an unsuccessful strategy, with the characterization of markers of different inheritance and evolutionary signal being preferred (COI, COII, ARN18S, ARN28S, ND6 and White gene) (Sallum, 2002; Bourke, 2011), to avoid genealogical discrepancies, ancestral polymorphisms and incomplete grouping of lineages, given that the subgenus Nyssorhynchus, is composed of 50% of species complexes, which has been considered a difficulty to infer evolutionary relationships, in addition to the use of previously characterized molecular markers in the identification of species to solve phylogenetic problems, demonstrating the low signal of ITS2 and COI-Barcode to infer ancestor-descendant relationships (Hoyos et al, 2015c; Hoyos et al. 2015d). In our inference, the individual genes show different degrees of resolution and phylogenetic signal, however, it was only the concatenated matrix under a partitioned scheme, the methodological treatment with higher resolution for the estimation with high supports of posterior probabilities of clade phylogenetic relationships between species of the subgenus *Nyssorhynchus*.

The phylogeny inferred from four mitochondrial and nuclear molecular markers clearly evidenced the paraphilia existing in the sections Myzorhynchella, Argyritarsis and Albimanus within the subgenus Nyssorhynchus. However, it is possible to highlight two defined groups: 1. Myzorhynchela sensu stricto (*An. lutzi + An. antunezi*) + Argyritarsis (*An. lanei, An. triannulatus* and *An. argyritarsis*) + Albitarsis Group (*An. albitarsis, An. deanorum* and *An. marajoara*). 2. Albimanus, consisting of the groups Strodei (*An. rondoni + An. strodei*) and Oswaldoi (*An. dunhami, An. goeldi, An. evansea, An. galvaoi, An. rangeli, An. konderi and An. oswaldoi*). Previous works failed to recover the relationships between the different groups of species representing the sections that correspond to the subgenus *Nyssorhynchus* (Sallum, 2000, Bourke, 2011, Foster, 2013), constituting the inferred phylogeny, the first hypothesis that allows to show clear relationships between members of this subgenre.

### Section Myzorhynchella

Foster et al (2013), find high levels of divergence for *An. parvus*. This species is not grouped with members of the same section (*An. lutzi and An. antunezi*) and is positioned as a sister group of all the species that contains the subgenus *Nyssorhynchus*. Sallum et al (2000), using morphological characters, showed similar relationships to those inferred by Foster et al (2013), and in both works the *An. parvus* elevation to the status of its own subgenus is proposed, differentiating it from the species that make up the subgenus *Nyssorhynchus*. The species *An. lutzii* and *An. antunezi* are highlighted as brother clade of the species *An. lanei, An. triannulatus* and *An. argyritarsis* of the Argyritarsis section.

### Section Argyritarsis

*An. darlingi* constitutes a sister group in ancestral position to the two inferred clades with the concatenated matrix. It is clear that this section should be reconsidered taxonomically, due to its evolutionary closeness with species of the Myzorhynchella section.

### Section Albimanus

Within this clade, *An. braziliensis* considered from the Argyritarsis section as a sister species of the Albimanus section is observed. This presents members considered mostly species complexes. The basal position is shared by *An. oswaldoi* and *An. konderi*, being the Oswaldoi group, the grouping that includes the most derived species within the subgenus *Nyssorhynchus*.

### Divergence estimates

Krzywinsky et al. (2011) estimated at 228 ma the separation of lineages corresponding to Anophelinae and Culicinae. Reidenbach et al. (2009) inferred with six nuclear genes in 51 ma, the age of separation of the genera *Anopheles* and *Bironella*, considering that the branch of the genus *Anopheles*, made up of species from the subgenres *Stehomyia* and *Anopheles*, has an age of 43 ma, agreeing with its with the estimated age for the fossil *An*. (*Nyssorhynchus*) *dominicanus* (Zavortink, 2000) by Cepek (cited by Schleek, 1990) for the strata where the amber of this species was found (30 – 45 ma). In our chronogram, the age of the *Nyssorhynchus* lineage is in the interval 27.2 – 24.01 ma, observing two times of diversification / speciation for the subgenus: 1). Species in ancestral branches diverge early from the separation of Nyssorhynchus from subgenus siblings (*Kerteszia* + *Anopheles*). 2). most of the species of the groups Albitarsis, Strodei and Oswaldoi present diversification mainly to associated events in the Pliocene (5.3 – 2.8 ma).

By way of conclusion, we can demonstrate that the treatment of molecular data prior to phylogenetic analysis and the characterization of molecular markers for evolutionary inference, can help to clarify the evolution of the subgenus *Nyssorhynchus* and answer questions about the evolution of vector competition for protozoa as *Plasmodium* spp. and the estimation of divergence times, allows the current understanding of the biogeographic patterns, and consequently may imply a better understanding of the epidemiological and evolutionary scenarios of malaria in South America. The current taxonomic hierarchy within the subgenus *Nyssorhynchus* should be reconsidered for the paraphilia between Myzorhynchella and Argyritarsis, and the subgenus elevation of the *An. parvus* complex. Also, the characterization of the markers used in this study to the remaining members of the subgenus *Nyssorhynchus* to determine the evolutionary affinities and have a holistic view of the taxonomy and evolution of the group.

